# Structural modeling of the SARS-CoV-2 Spike/human ACE2 complex interface can identify high-affinity variants associated with increased transmissibility

**DOI:** 10.1101/2021.03.22.436454

**Authors:** Hin Hark Gan, Alan Twaddle, Benoit Marchand, Kristin C. Gunsalus

## Abstract

The COVID-19 pandemic has triggered concerns about the emergence of more infectious and pathogenic viral strains. As a public health measure, efficient screening methods are needed to determine the functional effects of new sequence variants. Here we show that structural modeling of SARS-CoV-2 Spike protein binding to the human ACE2 receptor, the first step in host-cell entry, predicts many novel variant combinations with enhanced binding affinities. By focusing on natural variants at the Spike-hACE2 interface and assessing over 700 mutant complexes, our analysis reveals that high-affinity Spike mutations (including N440K, S443A, G476S, E484R, G502P) tend to cluster near known human ACE2 recognition sites (K31 and K353). These Spike regions are conformationally flexible, allowing certain mutations to optimize interface interaction energies. Although most human ACE2 variants tend to weaken binding affinity, they can interact with Spike mutations to generate high-affinity double mutant complexes, suggesting variation in individual susceptibility to infection. Applying structural analysis to highly transmissible variants, we find that circulating point mutations S447N, E484K and N501Y form high-affinity complexes (~40% more than wild-type). By combining predicted affinities and available antibody escape data, we show that fast-spreading viral variants exploit combinatorial mutations possessing both enhanced affinity and antibody resistance, including S447N/E484K, E484K/N501Y and K417T/E484K/N501Y. Thus, three-dimensional modeling of the Spike/hACE2 complex predicts changes in structure and binding affinity that correlate with transmissibility and therefore can help inform future intervention strategies.

## Introduction

The COVID-19 pandemic poses several interrelated scientific challenges that include understanding its biological origins and evolutionary trajectory, deciphering the mechanisms of virus-host interactions, and developing effective intervention strategies. Progress in understanding viral mutations will impact how we assess virus-host interactions and design intervention strategies. A major concern of the COVID-19 disease is that its causative virus SARS-CoV-2 has evolved to increase its ability to infect human hosts, as highlighted by recent identification of novel viral mutations in COVID-19 patients in multiple countries (cdc.gov/coronavirus/2019-ncov)^1^. Analysis of SARS-CoV-2 genome sequences suggests that there are thousands of mutations whose biological and health implications are unknown (CoV-GLUE database). It is thus important to develop methods that can effectively identify biologically significant COVID-19 mutations which can serve as surveillance tools for emerging infectious strains and guide development of new vaccines.

Advances in understanding the molecular basis of SARS-CoV infection in the last 15 years have led to the identification of mutations that enable coronaviruses to jump from animal to human hosts, as well as the molecular mechanisms underlying viral entry into human cells^2; 3; 4; 5^. Coronaviruses, including SARS-CoV, SARS-CoV-2 and MERS-CoV, are enveloped viruses with a positive-sense, single-stranded RNA genome. Several structural proteins are embedded in the viral envelope, notably the Spike (S) proteins, which are heavily glycosylated and assemble into homotrimer complexes that bind human cell surface receptors and thereby help the virus gain entry into its host^5; 6^. The S protein is composed of S1 and S2 subunits, which respectively form, the head and stem of the protein. The S1 subunit contains the host receptor-binding domain (RBD, residues 318-510 for SARS-CoV); the S2 subunit is membrane-associated and anchors the Spike protein in the viral envelope. Coronaviruses SARS-CoV, SARS-CoV-2 and RaTG13 use the S RBD to enter host cells by binding to human angiotensin-converting enzyme 2 (hACE2) receptors, triggering the subsequent virus-host cell fusion event^4; 5; 6; 7; 8^. This central mechanism of SARS-CoV and SARS-CoV-2 infection accounts for the importance of the S protein in understanding cross-species adaptation^4^ and current therapeutic development^9; 10; 11; 12^. In contrast to SARS-related viruses, the Spike proteins of MERS-CoV and HCoV-229E coronaviruses use the dipeptidyl-peptidase 4 (DPP4) and aminopeptidase N (APN) receptors, respectively, to gain entry into host cells^5^.

Recognition of the ACE2 receptor by S RBD is a limiting step in viral infection and cross-species adaptation in SARS-CoV strains^4; 13^. Biochemical and crystallographic studies of SARS-CoV S-ACE2 complexes have identified key amino acid residues responsible for binding specificity (Figure 1a). Specifically, the determinants of high-affinity ACE2 association are the interaction interface residues N479 and T487 in the SARS-CoV S-protein RBD^2^. On the complementary ACE2 interaction surface, residues K31 and K353 are sensitively influenced by S-protein positions 479 and 487, respectively. Consequently, residues K31 and K353 and their vicinities define the hACE2 interaction hotspots for SARS-CoV S proteins isolated from animals and humans^2; 14^.

**Figure 1.**
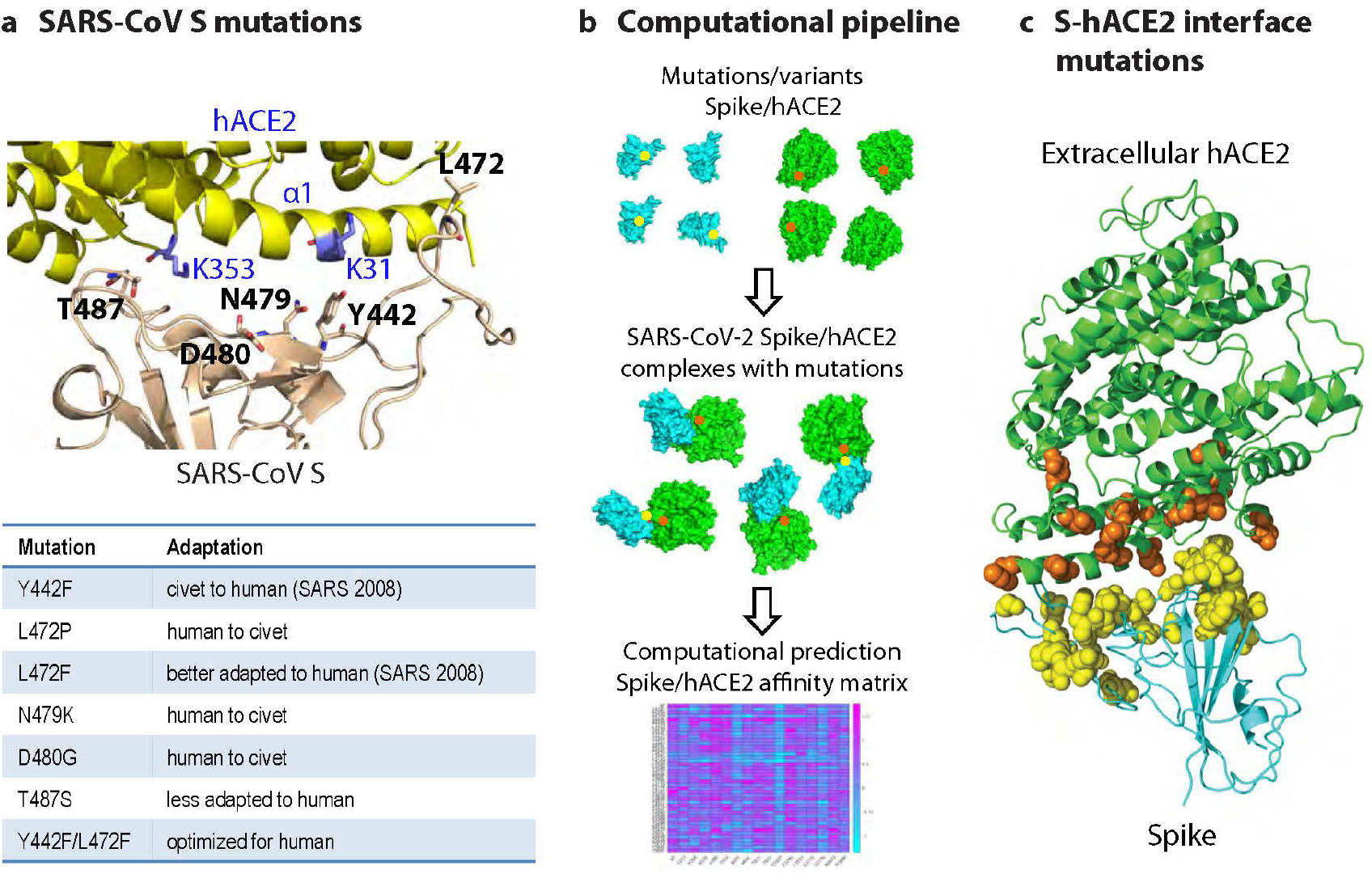
Computational framework for predicting the effects of variants in the SARS-CoV-2 Spike/human ACE2 complex. (**a**) Top: SARS-CoV S-hACE2 complex showing key S interface residues and mutations that play a role in cross-species adaptation to hACE2 (listed in table below). Recognition of hACE2 by the S protein is centered at hACE2 residues K31 in the α1 helix and K353 (highlighted in blue). Bottom: Known adaptations of selected S protein mutations. (**b**) Computational workflow for structural assessment of variants in the S-hACE2 complex. First, genetic variants are identified bioinformatically from publicly available genome sequence data. Second, starting from their native conformations, mutations in 3D structures of component proteins are generated and the mutant complex is then assembled and refined using an energy minimization algorithm. Third, the binding affinity of each mutant S-hACE2 complex is computed using physics-based interaction energies, including solvation, van der Waals, electrostatics with ionic effects, and entropy. (**c**) Model of a wild-type (WT) SARS-CoV-2 S-hACE2 complex highlighting positions of modeled variants at the interface (spheres). Spike residues, yellow; hACE2 residues, orange.

Analysis of SARS-CoV cross-species adaptations has uncovered specific residues in the S-protein that are favored by ACE2 from different species (Figure 1a)^15^. For example, hACE2 prefers residue N at S-protein position 479, whereas K, N and R residues are found in palm civet ACE2. At S-protein position 487, both human and civet ACE2 favor residue T, but residue S is much less preferred in humans. Other S-protein determinants of ACE2 association and cross-species adaptation include residues Y442, L472, and D480^14^. Although Y442 is found in civet and human SARS-CoV, F442 is thought to be preferred in human hosts, as found in the SARS-CoV strain isolated in 2008. At position 472, F is adapted to hACE2, whereas P to civet; at position 480, hACE2 favors residue D, whereas civet ACE2 prefers the G residue. Interestingly, enhanced adaptation to hACE2 can be engineered by simultaneous Y442F and L472F mutations^14^. Thus, residue-level analysis sheds light on the vital role of S RBD mutations in receptor recognition, viral transmission and cross-species infection. Since structural analyses have revealed extensive similarities between the receptor binding modes of SARS-related viruses, our accumulated knowledge of SARS-CoV will provide an indispensable guide to analysis of mutations in the SARS-CoV-2 S-hACE2 receptor complex^6; 8; 16; 17; 18^.

Single mutations in ACE2 can also have significant impact on host susceptibility to infection. Rats are resistant to SARS-CoV infection largely because of a single residue difference from hACE2 at position 353, one of its interaction hotspots. By mutating residue K353 to H found in rats, the variant human ACE2 has a low affinity for the S protein^2^. Likewise, for other species, residue differences in the ACE2 interaction interface lead to varying affinities and susceptibilities to SARS-CoV^13^. Moreover, analysis of hACE2 mutations using deep mutagenesis has provided insights into the specificity of SARS-CoV-2 S-hACE2 interactions and development of engineered soluble receptors as S decoys^19^. Thus, a complementary assessment of the role of hACE2 variants is needed to provide a comprehensive map of SARS-CoV-2 S-hACE2 interactions. Global surveillance of SARS-CoV-2 using genome sequencing has identified over 5000 amino acid substitutions, deletions and insertions in the S protein alone (CoV-GLUE). This large pool of changes poses a significant challenge for functional screening, which will require multi-level analysis encompassing sequence, structural, cellular and whole animal approaches. Recent emergence of fast-spreading SARS-CoV-2 lineages in UK, South Africa, Brazil and other countries reinforces the need for surveillance and to anticipate infectious mutations (nextstrain.org/sars-cov-2). Specifically, the recurring S mutations E484K, N501Y, D614G and their combinations have been implicated in increased viral transmission. The widespread S protein mutation, D614G, has also been found to increase viral infectivity in cell cultures^1; 20; 21^. Structural modeling suggests that D614G promotes an open conformational state, making the S-protein available for receptor binding^22^. Thus, concerted efforts to structurally and functionally characterize the new variants are urgently needed to uncover their contributions to receptor recognition and impact on the efficacy of current therapeutics and vaccines.

Here we apply a structural modeling approach to enable efficient screening of many mutant SARS-CoV-2 S-hACE2 complexes by computing their binding affinities, which have been shown to be a reliable measure of SARS-CoV’s potential to infect host cells^2; 3; 4^. Our approach, originally developed for large-scale modeling of microRNA-protein complexes^23; 24; 25; 26^, aims to fill the gap between sequence-based analysis and laboratory confirmation of functional mutations, which is often time consuming and resource-intensive. Importantly, we show that our method reproduces known affinity trends associated with key SARS-CoV mutations in cross-species adaptation mutations identified in previous SARS-CoV studies, as well as infectivity data for SARS-CoV-2 S mutations. We then assess the effects of 731 natural variant combinations at the interface of SARS-CoV-2 S and hACE2 proteins. We found that 31 high-affinity S mutations, including N440K, S443A, G476S, E484R, G502P, are clustered in the vicinities of two known interaction hotspots of hACE2. Significantly, we show that computed binding affinities for recent fast-spreading variants have high affinities (in decreasing order of affinity, S477N/E484K, E484K/N501Y, K417T/E484K/N501Y) and are resistant to neutralizing antibodies or overlap with their binding sites, consistent with their rapid spread and emergence of combinatorial mutations. Together these results suggest productive regions for future experimental work and indicate that structural modeling can be an effective surveillance tool for tracking emerging mutations in infectious COVID-19 strains.

## Results

### Structural modeling captures key aspects of Spike-hACE2 interactions

To ascertain the accuracy of our structural modeling approach (Figure 1b), we compared measured and predicted affinities of multiple SARS-CoV and SARS-CoV-2 variants for hACE2. We modeled the S-ACE2 complex as the interaction between S RBD (residues ~320 to ~520) and the extracellular N-terminal peptidase domain of ACE2 (residues ~20 to 613). Experimental affinity measurements using bio-layer interferometry and surface plasmon resonance (SPR) have shown that hACE2 binds to SARS-CoV-2 S-RBD with a greater affinity than to SARS-CoV S-RBD (Table 1). Specifically, the measured ratio K_D_(SARS-CoV)/ K_D_(SARS-CoV-2) is greater than 4 for both experimental methods. Our computed binding affinities confirm that hACE2 has a stronger affinity for SARS-CoV-2 S-RBD than for SARS-CoV S-RBD (−41.1 kcal/mol versus −22.0 kcal/mol, Table 1). For both native complexes, the favorable solvation (hydrophobic) and van der Waals interactions are balanced by unfavorable electrostatic and entropic forces.

**Table 1.** Measured dissociation constants from different experiments and predicted binding free energies from structural modeling. S1, S^B^ and S-CTD contain S-RBD.

| Viral protein domain | Method | K<sub>d</sub> (nM) | ΔG (kcal/mol) | Reference |
| --- | --- | --- | --- | --- |
| SARS-CoV S<sup>B</sup> | Biolayer interferometry | 5.0 ± 0.1 |  | Walls, AC et al 2020 |
| SARS-CoV S-RBD | SPR assay | 408.7 ± 11.1 |  | Wang, Q et al 2020 |
| SARS-CoV S-RBD | SPR assay | 325.8 |  | Wrapp, D et al 2020 |
| SARS-CoV S-RBD | Structural modeling |  | -22.0* | This work |
| SARS-CoV-2 S<sup>B</sup> | Biolayer interferometry | 1.2 ± 0.1 |  | Walls, AC et al 2020 |
| SARS-CoV-2 S1 | SPR assay | 94.6 ± 6.5 |  | Wang, Q et al 2020 |
| SARS-CoV-2 S-CTD | SPR assay | 133.3 ± 5.6 |  | Wang, Q et al 2020 |
| SARS-CoV-2 S-RBD | SPR assay | 14.7 |  | Wrapp, D et al 2020 |
| SARS-CoV-2 S-RBD | Structural modeling |  | -41.1 | This work |

\* Average of two native complexes

Since certain mutations at five positions (Y442, L472, N479, D480, and T487) are responsible for the ability of SARS-CoV to adapt to human from animal ACE2 (Figures 1a)^4; 14; 15^, a sensitive test of our structural modeling approach is whether it can reproduce the expected binding affinity patterns for such mutations. We consider seven cases representing distinct animal-human adaptive mutations in the SARS-CoV S-protein (Figure 2a)^3^: adaptations from civet, an intermediate animal host, to human ACE2 (Y442F, L472F); mutations obtained by reversing civet to human adaptations (L472P, N479K, D480G); unfavorable mutation (T487S); and a previously designed double mutant (Y442F/L472F) optimized for hACE2 binding. A plot of the computed affinities versus measured association constants^3^, 1/ K_D_, shows that structural modeling captures the overall trend in measured affinities produced by the key SARS-CoV S protein mutations (R^2^=0.61; Figure 2a). Examining specific examples, the point mutations Y442F and L472F, which are adapted to hACE2, produced better affinities relative to the WT complex (−26.0 kcal/mol and −27.8 kcal/mol versus −22.0 kcal/mol). Interestingly, the optimized Y442F/472F double mutant generated a very high affinity complex (−42.6 kcal/mol), consistent with association (1/K_D_ value over 3 times greater than that for L472F) and infectivity data^3^. When two of the hACE2-adapted mutations in SARS-CoV are reversed to those found in civet (L472P, N479K), the predicted affinities decreased dramatically (−4.9 kcal/mol and −14.8 kcal/mol, respectively). Lastly, the mutation T487S, which is judged to be less adapted to human ACE2 based on frequencies of occurrence of these mutants in civet and human SARS-CoV, yielded a poorer affinity (−16.5 kcal/mol), in agreement with expectation.

**Figure 2.**
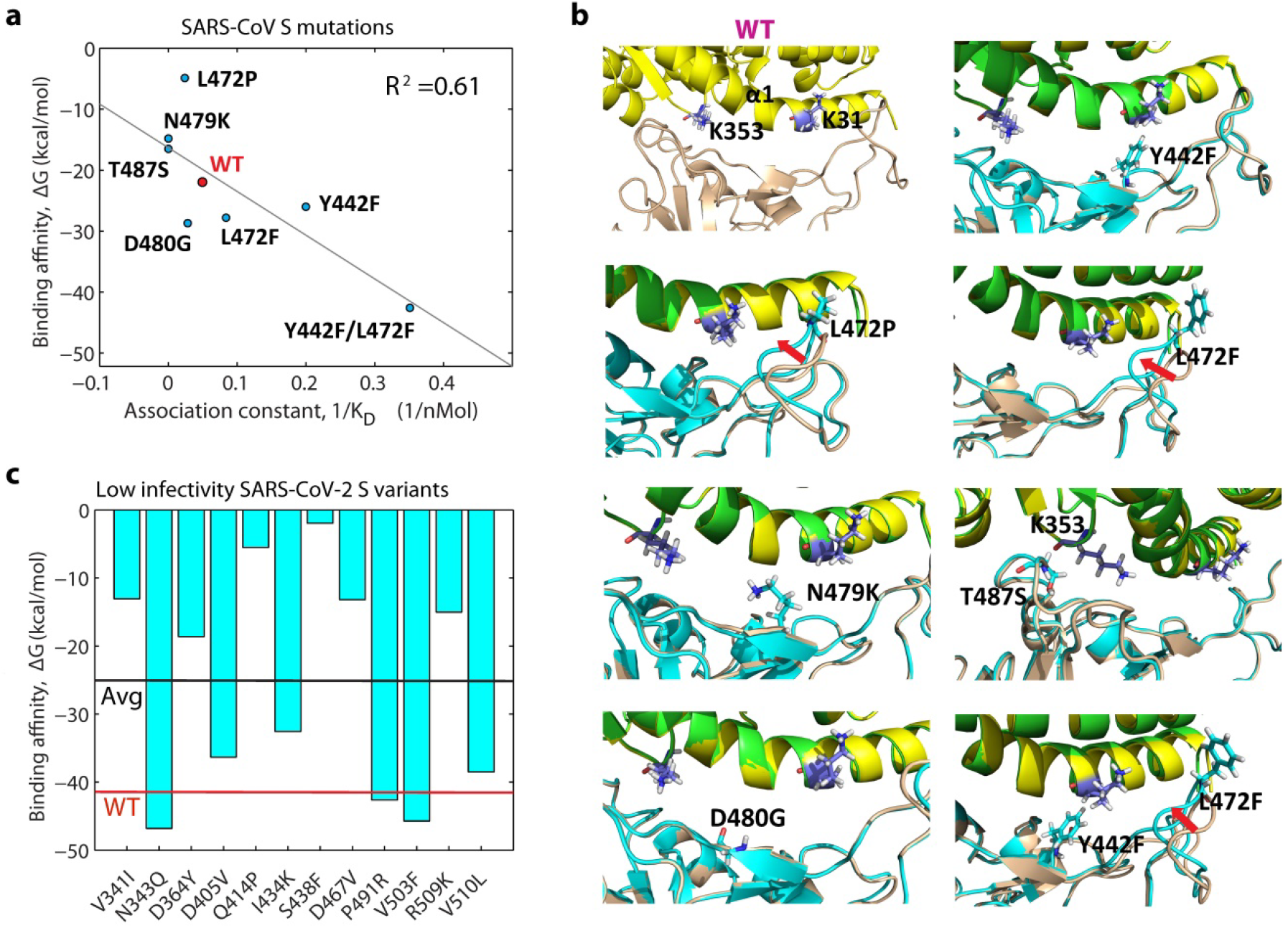
Comparison of measured and predicted affinities of Spike mutations. (**a**) Comparison of measured association constants and predicted affinities of seven key SARS-CoV S mutations and WT complex (PDB:2AJF, upper left panel); the fitted regression line has R^2^=0.61. (**b**) Conformational changes induced by SARS-CoV S mutations shown in (a). Mutations L472F, L472P and Y442F/L472F produced local conformational distortions at the interaction interface (S-cyan, hACE2-green), whereas mutations Y442, N479K, D480G, and T487S do not structurally perturb the wild-type complex (S, wheat; hACE2, yellow). Red arrows indicate structural movements. (**c**) Predicted affinities of natural SARS-CoV-2 S variants that have low measured infectivity levels relative to WT have significantly lower predicted affinities: −25.8 kcal/mol on average (black line) versus −41.1 kcal/mol for WT (red line).

Another test of our method is to compare predicted binding affinities to measured infectivity data for variants in the SARS-CoV-2 S-hACE2 complex. Quantification of the effects of S variants using infectivity assays is correlated with their binding affinities, as well as other factors (dynamics of S and its interactions with other host proteins). A recent work identified 12 natural variants in the SARS-CoV-2 S-protein at the interface that show significantly lower infectivity (based on luciferase reporter levels) in comparison with the highest-frequency variant, which we call “wild type (WT)”^27^. Predicted affinities for these mutations indicate that 9 out of 12 cases have lower binding affinities compared to WT, and the mean affinity of all cases is reduced by 40% (−25.8 kcal/mol compare to −41.1 kcal/mol for WT; Figure 2b). Thus, structural modeling correctly reproduces affinity trends for previously characterized SARS-CoV and SARS-CoV-2 S mutations, suggesting that the approach is sufficiently accurate for general investigations of SARS-CoV-2 S-hACE2 mutant complexes.

### High-affinity SARS-CoV-2 Spike mutations are clustered near hACE2 interaction hotspots

Using 16,083 sequenced genomes (downloaded May 4, 2020), we found 739 SARS-CoV-2 S single amino acid replacements based on our variant calling pipeline (Suppl. Figure 1). We chose 42 S and 16 hACE2 interface variants to produce 731 possible combinations of complexes with zero or one mutation in each protein (Methods). We represent associations for all 731 mutant complexes as a normalized affinity matrix, ΔG_mutant_/ΔG_WT_ (heatmap in Figure 3a). There are 31 high-affinity complexes, defined as >20% of ΔG_WT_ (magenta), 111 complexes with WT-like affinities (±20% of ΔG _WT_), 435 low-affinity complexes (20% below ΔG _WT_ but <0), and 154 complexes with unfavorable formation energies that are predicted to remain unbound (ΔG_mutant_ >0, blue). Thus, mutations that produce high affinity complexes are relatively rare (4%), while low-affinity mutations are relatively common (60%). The unbound complexes (ΔG_mutant_ >0) can be interpreted to mean that some S-hACE2 mutation combinations are incompatible with each other and therefore most likely do not occur in COVID19 patients.

**Figure 3.**
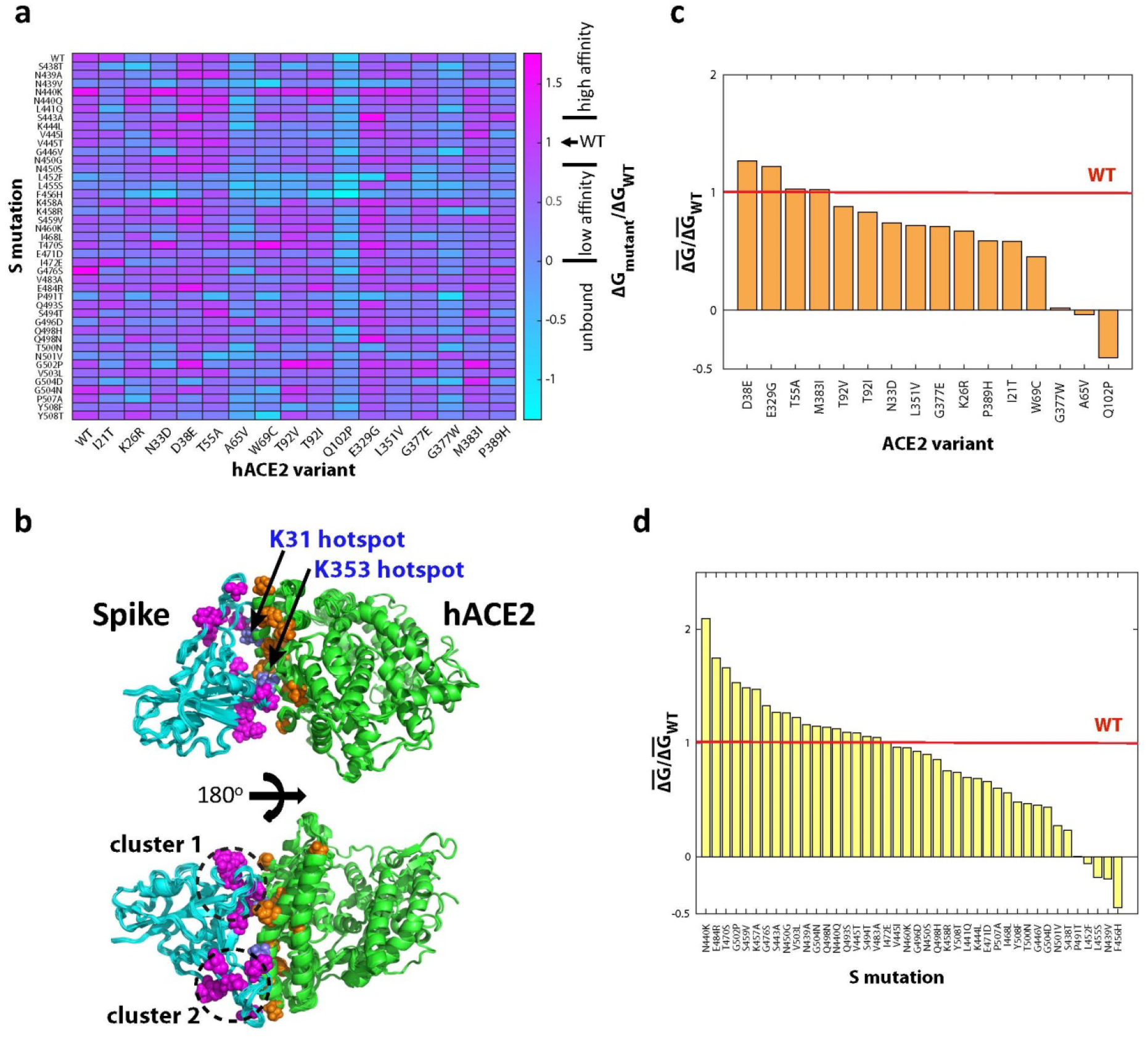
Affinities of interface variants in the SARS-CoV-2 Spike/human ACE2 complex. (**a**) Binding affinity matrix for SARS-CoV-2 S-hACE2 complexes, representing all pairwise combinations of 16 hACE2 variants and 42 S mutations. Each component protein has either zero or one mutation. The affinity heatmap represents affinities normalized by that of the WT complex (PDB: 6M17). Normalized affinities (relative to WT) are classified into four categories: high-affinity (>1.2), WT-like (1±0.2), low-affinity (<0.8 but >0), and unbound (<0). (**b**) Superimpositions of 31 high-affinity mutant S-hACE2 complexes. Highlighted are the associated mutations in S (magenta spheres) and hACE2 (orange spheres), as well as hACE2 interaction “hotspots” at K31 and K353 (blue spheres). On S protein, the high-affinity mutations form two clusters at opposite ends of its binding interface (dashed circles); 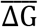 and 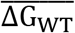 are mean affinities of mutant and WT complexes, respectively. (**c**) Normalized mean affinities of hACE2 variants averaged over all S mutations. (**d**) Normalized mean affinities of S mutations averaged over all hACE2 variants.

Superimpositions of all 31 high-affinity complexes show that their S mutations form two clusters at opposite ends of S RBD’s concave interaction surface (Figure 3b, magenta spheres). Solvent-exposed ends of the receptor binding motif (RBM, residues 440-505), a large loop that directly binds hACE2, are more susceptible to conformational fluctuations; thus, the clustering of mutations in these regions likely reflects the potential to generate more stable or higher affinity complexes. An exception to this rule is the high-affinity mutant S494T, which occurs near the short beta strand in the center of S interaction surface. The rare occurrence of high-affinity mutations in the central RBM interface region could imply that the opposite-facing hACE2 α1 ridge region does not tolerate S mutations. Overall, our identification of high-affinity S mutations suggests structural regions at the interface where they are most likely to occur.

### Complexes with a single mutation in either the Spike or hACE2 protein

To focus on more common complexes in infected individuals, we analyzed the affinities of complexes in which one of the proteins represents the WT (highest frequency) genetic variant (Figure 3a, first row and column). Related experimental studies have focused on random mutations in either S or hACE2, generated using error-prone polymerases^19; 28^. Narrowing our focus to naturally occurring variants at the interaction interface, we find that structural modeling of hACE2 variants generally result in lower affinity, whereas Spike variants span a wide affinity range.

We found no high-affinity hACE2 variants (20% above WT affinity) and, with the exception of D38E and E329G, most hACE2 variants in fact lead to lower binding affinities (Figure 3a). These results imply that with WT S as the binding partner, most hACE2 variants reduce the ability of the SARS-CoV-2 virus to enter host cells. This is consistent with experimental assessment of 165 natural hACE2 variants from mutagenesis experiments, which showed that only 5% of hACE2 variants have increased affinity for WT S, whereas 65% have decreased affinity and 27% have similar affinity as WT hACE2^29^.

Two Spike variants (N440K and G476S), both in solvent-exposed residues at opposite ends of the binding interface, form high-affinity complexes in combination with WT hACE2 (Figure 4a,b). Compared with the WT complex, N440K gains 6 kcal/mol in solvation energy and 18 kcal/mol in van der Waals energy, whereas G476S gains 8 kcal/mol in van der Waals energy and 27 kcal/mol in electrostatic energy. These high-affinity cases indicate that improved affinities are produced by favorable short-range (van der Waals) and environment-specific (solvation and electrostatic) interactions.

**Figure 4.**
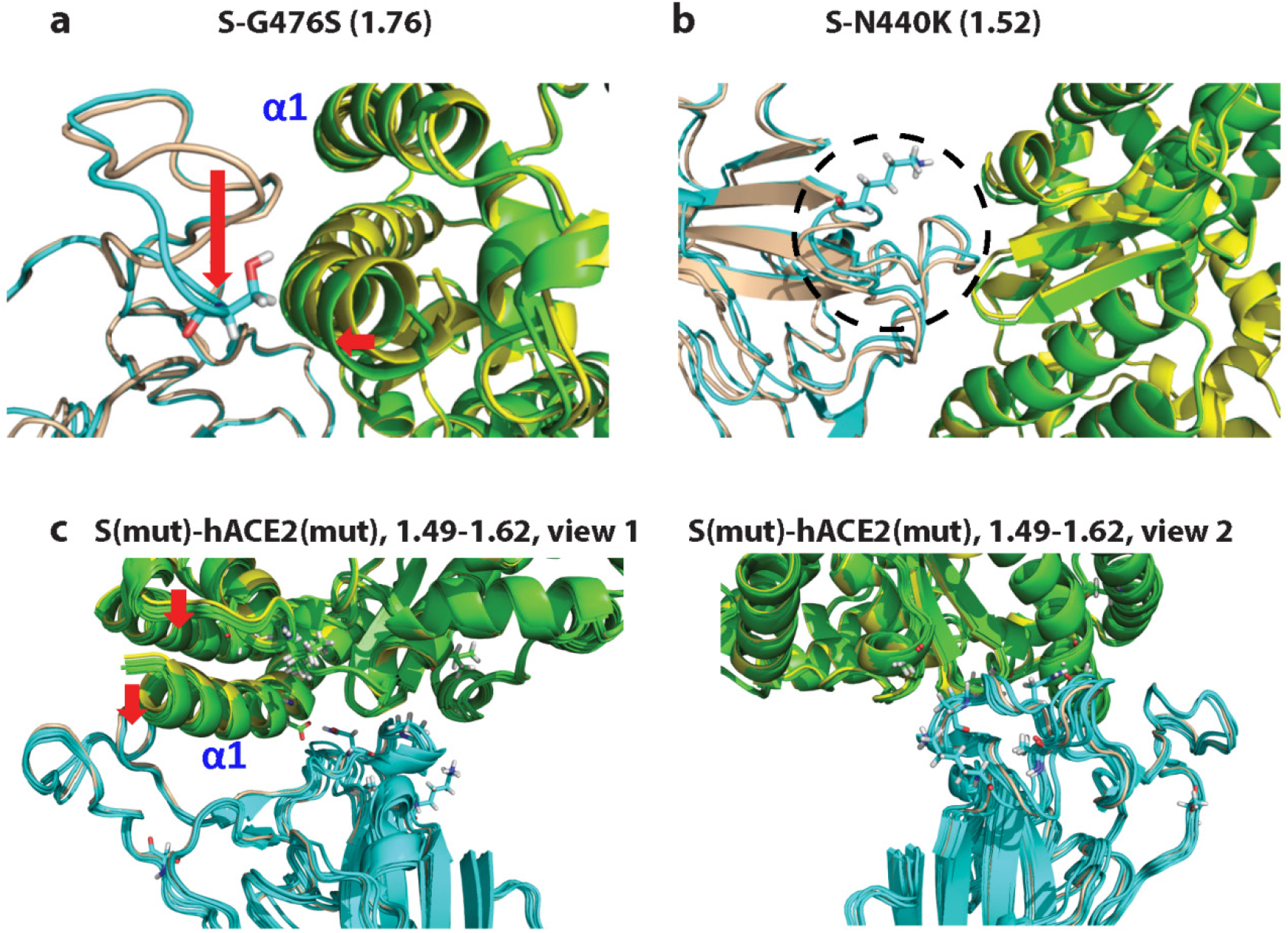
Conformational changes induced by high-affinity mutations. Local conformational distortions induced by single Spike mutations N440K (**a**) and G476S (**b**) and their associated normalized affinities. (**c**) Superimpositions of seven complexes with a mutation in each protein, S(mut)-hACE2(mut) and the range of their normalized affinities. These double-mutant complexes induced conformational changes (red arrows) in hACE2 α1 (labeled in blue font) and loops in S (437-452, 494-507) to yield enhanced binding affinities. Color scheme: WT complex, hACE2 in yellow and S in wheat; mutant complexes, hACE2 in green and S in cyan.

Among the unfavorable Spike mutations, L452F and P491T occur at or near the antiparallel beta strands located in the middle of S-RBM’s concave interaction surface, supporting our observation about the rarity of high-affinity mutations in this region. Compared with the WT complex, these cases have unfavorable electrostatic interactions (loss of ~30 kcal/mol) and an increase in entropic energy (~10 kcal/mol). Thus, the effects of S mutations vary across its binding interface and are influenced by the properties of the opposite-facing hACE2 binding surface. Even though some S mutations are unlikely to form complexes with WT hACE2, they can interact favorably with specific hACE2 variants (discussed below), suggesting that compensatory changes in the partner protein can promote binding.

### Complexes with a single mutation each in Spike and hACE2 proteins

Experimental studies of mutations in the S-hACE2 complex have focused on amino acid changes in either Spike or hACE2 protein^19; 28^. This limitation is due to the geometric increase of combinatorial space when mutations in both proteins are considered. Considering only complexes with mutations at the binding interface (Figure 3a, excluding first row and column), we find that complexes with a mutation in both proteins account for 29 of the 31 high-affinity complexes we identified. This suggests there is a nontrivial probability of generating mutually-reinforcing mutations in the S-hACE2 complex. Indeed, the hACE2 variants N33D, D38E, T92V, E329G and M383I interact favorably with many S mutations (Figure 3a). The top 7 high-affinity double mutation cases, in decreasing affinity, are S-S443A/hACE-E329G, S-G502P/hACE2-T92V, S-T470S/hACE2-W69C, S-N440K/hACE2-T92I, S-S443A/hACE2-D38E, S-Q498N/hACE2-E329G, and S-G502P/hACE2-M383I (superimposed and individual complexes in Figures 4c and Suppl. Figure 3). In contrast, three hACE2 variants (A65V, A102P and G377W) interact unfavorably with most S mutations examined. Using binding affinity as a proxy for viral infectivity at the cellular level, these results indicate that the patterns of favorable and unfavorable combinations represented in the affinity matrix can be exploited as a map to predict susceptibility of individuals or groups with specific hACE2 variants to infection by viral strains with mutations in the Spike protein.

### Conformational changes induced by high-affinity mutations in the Spike-ACE2 complex

To probe the observed mutation patterns more deeply, we examined the conformational changes induced by high-affinity mutations in the S-hACE2 complex. For this purpose, we considered the top 9 high-affinity complexes (Figure 4 and Suppl. Figure 3). Relative to the WT complex (Spike, wheat; hACE2, yellow), the single Spike G476S mutation generated a large conformational switch in a local RBM loop (residues 475-AGST-478) causing it to orient toward the N-terminal of hACE2 α1 helix, a region found to have more contacts with the S-protein of SARS-CoV-2 than with SARS-CoV (Figure 4a)^16^. This switch dramatically improved the electrostatic energy of the complex (from 18.2 kcal/mol to −9.0 kcal/mol); the short-range van der Waals energy also improved by 9 kcal/mol. The single Spike N440K mutation occurs near the anti-parallel β-sheet core of RBD but produced an extended perturbation in the RBM loop region facing the hACE2 K353 hotspot (Figure 4b). This mutation improved the solvation (by 6.3 kcal/mol) and van der Waals (18.0 kcal/mol) energy contributions to the binding affinity.

Turning to high-affinity, double mutant complexes (Figure 4c and Suppl. Figure 3), most have improved contact van der Waals energies (gain of 10-20 kcal/mol), and complexes S-G502P/hACE2-T92V and S-T470S/hACE2-W69C also have enhanced electrostatic energies (gain of ~30 kcal/mol). Superimpositions of these complexes reveal the structural movements accounting for gains in their energy components (Figure 4c). On the hACE2 interface, variants T92V/I cause the α1 helix and a loop (residues 87-92) to move closer to the S interface, reflecting improved van der Waals energies. On the S interface, four mutations (N440K, S443A, Q498N, G502P) generated significant conformational changes in two loops (437-452, 494-507). In conclusion, high-affinity mutations in S and hACE2 proteins produce structural changes in both proteins to optimize the interaction interface.

### High-affinity Spike mutations with the potential to increase transmission efficiency

Multiple lines of evidence can be used to suggest the most likely candidates that can potentially increase SARS-CoV-2 infectivity. To derive high-confidence predictions, we consider the common S variants from the 10 top-ranked affinities both among all individual variant combinations and averaged over all hACE2 variants (Figure 3a,d). This criterion ensures consensus between the two ways (individual and averaged affinities) of assessing high-affinity complexes. The SARS-CoV-2 S mutations fulfilling this stringent criterion are N440K, S443A, G476S, E484R and G502P. The affinity ranking of G476S is dominated by its binding to WT hACE2, whereas the other three S mutants interact favorably with multiple hACE2 variants (Figure 3a).

The significance of the high-confidence, high-affinity S variants can be analyzed based on sequence conservation of their residue positions and their proximity to the key residues in SARS-CoV. Multiple sequence alignment of SARS-CoV strains (from human, civet, and bat) and SARS-CoV-2 implies that the equivalent receptor recognition residues for SARS-CoV-2 S are L455, F486, Q493, S494, and N501^15^. Although the predicted high-affinity S variants do not overlap with these positions, sequence alignment shows that SARS-CoV-2 S N440 and G502 are 100% conserved and its S443, G476 and E484 positions are conserved in SARS-CoV strains, suggesting that these interface residues must play a role in receptor recognition and that certain mutations at these positions optimize virus-host interactions. In particular, at position 443, the corresponding residue in SARS-CoV strains is a conserved A, so the S443A mutation in SARS-CoV-2 could account for its enhanced binding to hACE2.

The high-confidence mutations are also predicted to alter the interaction interface (Figure 4 and Suppl. Figure 3). The G502P mutation, which is adjacent to the key N501 residue, alters the local interface conformation, providing structural support for increased S-hACE2 binding (Figure 4a). The N440K mutation involves a charge change and an extended conformation alteration in the RBM loop (Figure 4b). Thus, combined conservation, energetic, and structural analysis implies that certain point mutations near the interface have the ability to reprogram the interface conformations to produce high-affinity S-hACE2 complexes, which is the basis for efficient transmission of SARS coronaviruses that utilize ACE2 for host entry.

### High-affinity Spike mutations N440K and E484K/R coincide with antibody escape mutants

Since solved complexes have shown that some promising neutralizing antibodies (NAbs) for clinical applications bind to the hACE2 interface region of the S protein^30^, it is interesting to determine whether our identified high-affinity variants (Figures 3 and 4) overlap with the binding sites of NAbs. A recent study has shown that certain S mutations are resistant to three NAbs (C121, C135 and C144) currently being evaluated for their therapeutic potential^28^. Among the antibody escape mutants identified using a cell-based infectivity assay, N440K is resistant to C135 and E484K to C121. Both of these mutations involve a net change in charge, a likely determinant of antibody resistance. By our affinity measures N440K is predicted both to be capable of escaping antibody C134 and to bind hACE2 with high affinity, attributes that make it a prime candidate for viral surveillance. If electrostatic energy is the deciding factor in C121 antibody resistance, then the E484K variant can be regarded as chemically similar to our top affinity candidate E484R; indeed, recent deep mutagenesis analysis showed that E484R is resistant to three neutralizing antibodies (COV2-2050,COV2-2096, COV2-2479; Figure 5b)^31^. Structural mapping of N440K and E484R variants confirms that they overlap, respectively, with the independent binding sites of C135 and C121 on the S protein (Figure 5a,b). Given the infectivity potential and resistance of N440K and E484K/R variants, there is a need to monitor their spread.

**Figure 5.**
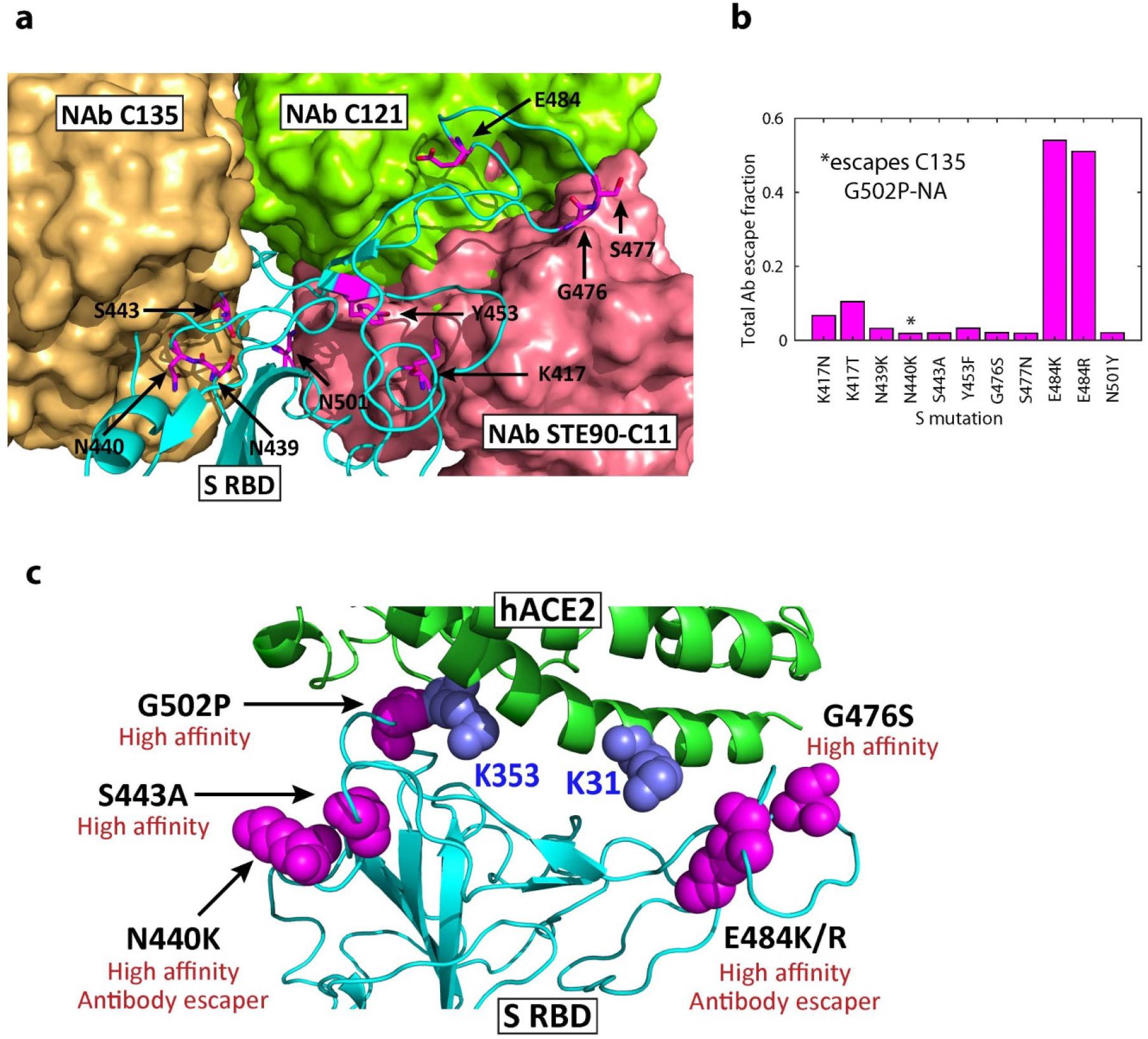
High-affinity mutations and antibody binding sites in Spike RBD. (**a**) Structure of WT S RBD superimposed with three neutralizing antibody structures (C121, 7K8X; C135, 7K8Z; STE90-C11, 7B3O). Residues predicted to harbor high-affinity variants and fast-spreading mutations are highlighted, illustrating that mutations in antibody binding sites have the potential to reduce their efficacy. Bound C135 and STE90-C11 antibody structures partially overlap. (**b**) Antibody escape of mutations as measured using the total escape fraction, i.e., the sum of all escape fractions across 10 antibodies tested using deep mutagenesis. (**c**) Summary of significant high-affinity mutations in S and hACE2 proteins and their functional implications.

### Structural/energetic basis of fast-spreading Spike variants

The newly identified S interface variants now in wide circulation across the globe include K417N/T, N439K, Y453F, S477N, E484K, N501Y and their combinations. To correlate affinity with transmission data, we consider the averaged affinity for all naturally occurring hACE2 variants, in order to account for their conformational repertoire and occurrence in different populations. Averaged affinities predicted by structural modeling indicate that the mutations fall into three groups (Figure 6a): very high-affinity (80 to 90% better that WT; S477N/E484K/N501Y, S477N/E484K and E484K/N501Y), high-affinity (40 to 50% better than WT; S477N, E484K, N501Y), WT-like (S477N/N501Y, K417T/E484K/N501Y, N439K, Y453F), and low-affinity (<70% of WT; K417T/E484K, K417T, K417T/N501Y).. The highest affinity group includes fast-spreading double mutants S477N/E484K and E484K/N501Y in Brazil, South Africa, US and UK (55587954 (medrxiv.org), cdc.gov/coronavirus/2019-ncov); the triple mutant S477N/E484K/N501Y with the best affinity has not been reported. The high affinity group consists of single mutants S477N, E484K and N501Y, indicating that the fast-spreading variants combine these mutations to achieve higher binding affinities; recent molecular dynamics simulations also predicted that S477N has a higher affinity than WT^32^. The WT-like group includes K417T/E484K/N501Y (spreading in South Africa, Brazil), N439K (Denmark) and Y453F (UK).

**Figure 6.**
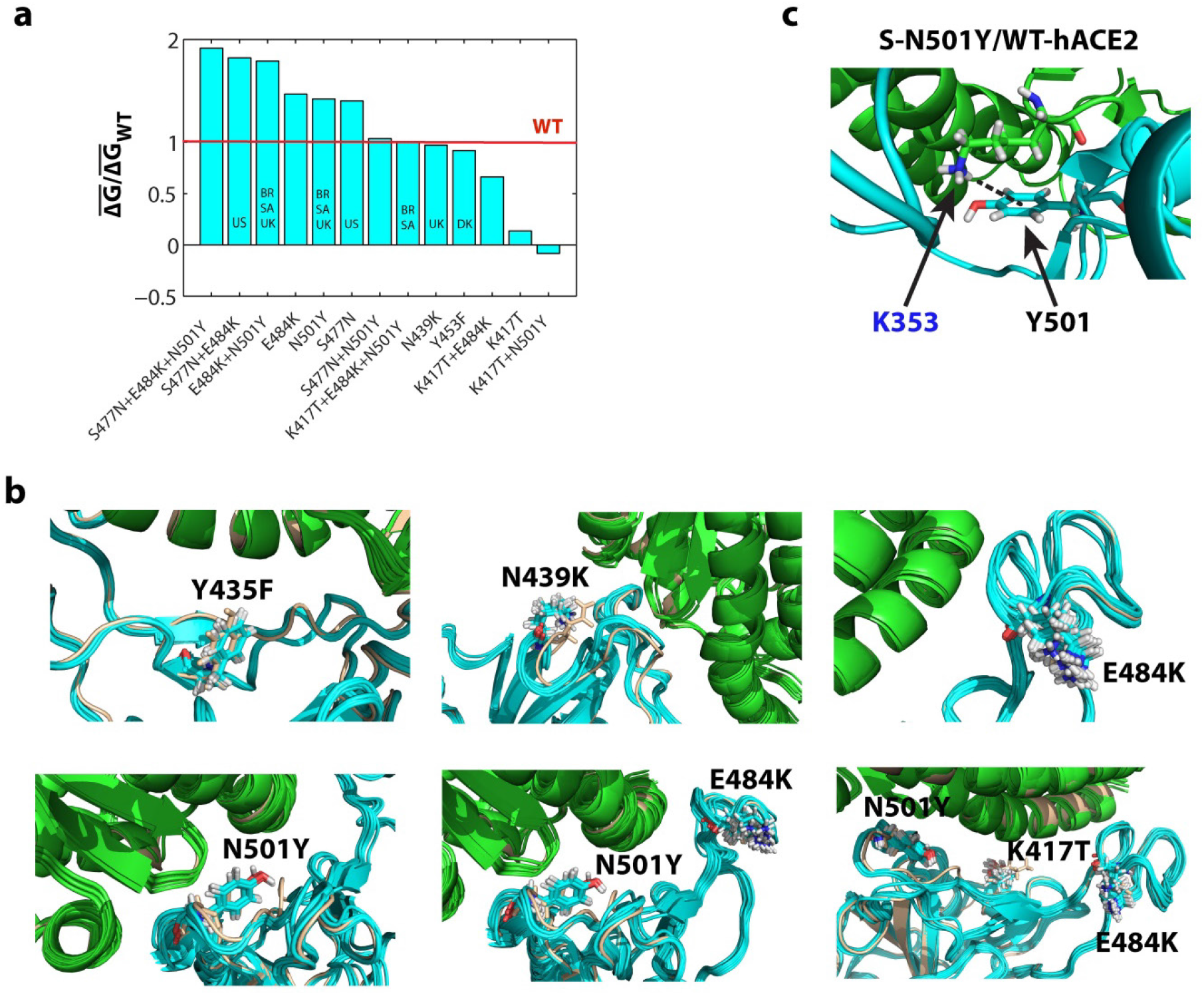
Structural and energetic analysis of fast-spreading Spike mutations. (**a**) Normalized mean affinities of mutations in the S-protein averaged over WT and 16 hACE2 variants. Prevalent variants circulating in different countries are indicated. Abbreviations: BR, Brazil; DK, Denmark; SA, South Africa; UK, United Kingdom; US, United States. E484K is present in the GSAID database; other unlabeled bars have not been observed in nature. (**b**) Structure ensembles of S-hACE2 complexes for each S mutation and the hACE2 variants. Each structure ensemble displays local conformational changes induced by a Spike mutation. (**c**) Configurations of Spike Y501 and hACE2 K353 sidechains suggest existence of cation-π interactions.

Antibody resistance is another crucial determinant of viral transmission.Mutants E484K/R exhibit high levels of antibody resistance (C121, COV2-2050, COV2-2096) and K417N/T have some degree of resistance (STE90-C11, COV2-2082), but the other mutations display negligible antibody resistance (Figure 5a,b). Thus, variants S477N/E484K, E484K/N501Y and K417T/E484K/N501Y indicate that they combine high affinity and antibody resistant mutations, which may account for their spread in multiple countries.

In contrast to E484K and N501Y, K417T alone appears not favored for receptor recognition, and this is reflected in reduced affinities for K417T/N501Y and K417T/E484K combinations, which are not observed together in sequenced viral genomes; K417N has similar characteristics (not shown). In vitro selection of optimized Spike RBD repeatedly recovered S477N, E484K and N501Y mutations but not K417N/T^33^. This indicates that K417N/T is not a major contributor to affinity, in agreement with our predicted binding affinities. Thus, the selective advantage of K417N/T is likely their ability to confer antibody resistance.

Superimposed structure ensembles for all hACE2 variants additionally reveal structural features that generate favorable complexes (Figure 6a). Although the E484K mutation does not interact directly with hACE2, it modifies the conformation of Spike RBD’s loop, resulting in a significant gain of average solvation energy (−11 kcal/mol). Further, K484’s sidechain shows considerable conformational variability in different hACE2 variant complexes (Figure 6b). In contrast, the mutant N501Y residue is closely packed in proximity of the K353 hotspot of hACE2 and its sidechain shows only minimal conformational changes. The favorable affinity for N501Y is produced by gains in entropic (6 kcal/mol) and van der Waals (7 kcal/mol) energies. Remarkably, the configurations of the lysine (hACE2 K353) and tyrosine (S Y501) sidechains suggest they could participate in favorable cation-π interactions, with the lysine being off-center of the tyrosine’s aromatic ring (Figure 6c). Estimation of the strength of this effect would require rigorous quantum-mechanical calculations. The high-affinity of the double mutant E484K/ N501Y is produced by gains in solvation (11 kcal/mol), van der Waals (2 kcal/mol) and entropic (5 kcal/mol) energies, indicating the approximate additive effect of its energy components. In sum, our analyses provide structural/energetic insights into the spread of the new SAR-CoV-2 variants.

## Discussion

Surveillance of biologically significant COVID-19 mutations is an important public health measure. While the implications of the vast majority of the ~5,000 S variants documented so far are unknown, already a handful of recently identified variants that appear to have increased transmissibility are spreading rapidly around the globe. As antiviral treatments and vaccines are being administered on a large scale, it is likely that selection pressures could alter viral evolutionary dynamics to favor resistant strains. Development of effective surveillance tools is needed to anticipate emerging mutations and inform treatment strategies.

Since detailed experimental analysis of infectivity for all possible combinations of hACE2 and S variants is impractical, we used in silico structural modeling of S-hACE2 complexes to predict structural changes and binding affinities for novel variants. Our exhaustive structural assessment of all combinations of naturally occurring mutations at the SARS-CoV-2 S-hACE2 interface has uncovered many mutants with significantly higher predicted affinities (>20%) relative to the WT complex. Surprisingly, many of these cases arise from mutually-reinforcing mutations in S and hACE2 proteins resulting from their favorable interactions. In biological terms, this means that carriers of certain hACE2 variants are predicted to be more susceptible to viral strains with specific mutations in the S protein. The high-confidence, high-affinity S mutations include N440K, S443A, G476S, E484R and G502P (Figure 5b), and among these N440K and E484K/R stand out for their ability to escape surveillance of some neutralizing antibodies (C135 and C121)^28^.

Our analysis of recently evolved, rapidly expanding strains with S mutations at the interaction interface provides insights into their relative affinities, molecular interactions, and the role of combinatorial mutations. We found that highly-transmissible S477N, E484K and N501Y mutations have greater average affinities than WT over known hACE2 variants. The double mutants S477N/E484K and E484K/N501Y have very high affinities (~80% above WT), which correlates well with its rapid spread in multiple countries (e.g., Brazil, South Africa, UK, US). Our structural analysis suggests that the widespread N501Y mutation likely introduces stabilizing cation-π interactions between S Y501 and hACE2 K353, whereas E484K enhances the solvation energy. Interestingly, our models predict that the triple mutation K417T/E484K/N501Y, which arose in the ancestral lineage E484K/N501Y, has about the same affinity as WT, although its frequency is increasing around the globe. The overlap of these positions with antibody binding sites confer robust resistance to neutralizing antibodies (Figure 5a,b)^28; 31^, thus offsetting the predicted loss of binding affinity to favor transmission of the triple mutant. Our analysis also predicts that the unobserved triple mutant S477N/E484K/N501Y has the highest affinity among all combinations of fast-spreading mutations examined.

Mutations outside of the interaction interface can also influence receptor recognition and viral transmission, as demonstrated by mutations identified in deep mutagenesis and in vitro evolution studies^19; 33; 34^. Indeed, identification of a stabilizing D614G mutation in the neck of the Spike protein that increases its density on viral particles^1^ highlights the need for further study of mutations outside of S RBD in the new SARS-CoV-2 lineages. Our predictive modeling of interface mutations in the S-hACE2 complex complements experimental studies^19; 34^ and supports the emerging view that combinatorial mutations in SARS-CoV-2 can simultaneously maintain high-affinity binding to hACE2 and evade antibodies (e.g., by N440K, E484K/R, K417N/T) in human hosts. We propose that structural modeling of novel variants as they arise could help predict which mutations may pose the greatest dangers in the ongoing pandemic and accelerate the development of new vaccines to protect against the spread of new highly infectious strains.

## Materials and Methods

### Computational pipeline

Since the S-hACE2 complexes for SARS-CoV^5; 35^ and SARS-CoV-2^6; 8; 16; 17^ have been solved, structural modeling provides a general method for analyzing mutations in these complexes. Based on previously established methodology, our computational pipeline for assessing mutations in complexes consists of the following steps (Figure 1b): (1) use bioinformatics algorithms to analyze SNPs and nonsynonymous mutations in proteins; (2) utilize solved protein structures to simulate mutations in component proteins, followed by structural refinement of the all-atom binary complex; and (3) compute the complex’s binding affinity score using physics-based interaction energies. Steps 2 and 3 are computationally intensive, requiring 3 days per complex on a linux cluster with parallel structure minimization runs.

### Bioinformatics analysis of SARS-CoV-2 Spike protein mutations

Mutations in the S protein were derived from SARS-CoV-2 genome sequences in GISAID, a publically available database. We used the original Wuhan SARS-CoV-2 sequence as the reference, wild-type sequence (NCBI’s NC_045512.2) for subsequent analysis. Genome data for submitted SARS-CoV-2 isolates were retrieved from GISAID on May 4, 2020, yielding a total of 16,083 sequences. To derive mutations in the S protein, we implemented the following computational pipeline (flowchart in Suppl. Figure 1). First, we aligned the reference S sequence to all annotated SARS-CoV-2 CDS sequences using promer, a script in the MUMMER4 package for rapid alignment of large DNA sequences. Second, from the aligned CDS sequences, we extracted SNPs using show-snps, a MUMMER4 utility program for reporting polymorphisms (SNPs) and insertions/deletions from promer’s alignment output. Third, we filtered for first Spike reading frame and removed ambiguous (e.g., nonsense) bases in SNPs. Finally, we determined the nonsynonymous variants and computed the frequency of each mutation. Our variant calling code is available on Github (https://github.com/GunsalusPiano/covid-snp-analysis).

### Variants in the human ACE2 protein

We used hACE2 variants identified in two previous studies^36; 37^. Since the variants that are most likely to influence Spike-hACE2 binding are those in the vicinity of the interaction interface, we narrowed the list to the following 16 hACE2 variants: I21T, K26R, N33D, D38E, T55A, A65V, W69C, T92V, T92I, Q102P, E329G, L351V, G377E, G377W, M383I, and P389H.

### Structural modeling of mutations in SARS-CoV Spike/hACE2 complexes

For SARS-CoV, we used the X-ray crystallographic complexes in PDB ID 2AJF (resolution of 2.90 Å) and for SARS-CoV-2, cryo-EM complexes in 6M17 (resolution of 2.90 Å). Based on these solved complexes, we modeled the S-hACE2 complex as the interaction between S RBD (residues ~320 to ~520) and the N-terminal peptidase domain of ACE2 hACE2 domain (residues ~20 to 613). We do not model the effects of N glycans at two S RBD modification sites (331 and 343), which are outside of the binding interface^38^, as well as the hACE2 modification site at N90.

From the mutation lists for Spike and hACE2 proteins described above, we focused on those mutations at the interaction interface since the effects of mutations are expected to diminish with distance from the interface. The focused mutation sets consist of 42 Spike mutations and 16 hACE2 variants (Suppl. Figure 2). We generated point mutations using the mutate.pl program in mmtsb package developed by Charles Brooks III’s laboratory. For each protein, the algorithm rebuilds the structure for the new sequence using the original C_α_ and side chain center positions while preserving the coordinates of unmutated residues. To avoid perturbations introduced by structure refinement, we generated all mutations using the experimentally solved structures. The reassembled binary complexes with mutations were then refined using a local minimization (L-BFGS) algorithm as implemented in the Tinker package. To ensure convergence, we use a cutoff rms gradient of 0.01 kcal/mol/Å or a maximum of 6000 iterations.

The binding free energy of a binary complex (ΔG) is expressed as the difference between the binding energies of the complex and its unbound components: ΔG = G_Spike-hACE2_ - G_Spike_ - G_hACE2_. This formula implies computing ΔG requires minimization of the binary complex and its unbound components. To efficiently run hundreds of complexes, the minimizations were computed in parallel. Computationally, ΔG, which is a function temperature (T) and ion concentration (c_ion_), is decomposed into different interaction types, ΔG(T,c_ion_) = ΔG_solvation_ + ΔG_vdw_ + ΔG_electrostatics_ + ΔG_entropy_. The ionic effects were computed using the Poisson-Boltzmann equation solver APBS at c_ion_=150mM NaCl and T=37°C. Electrostatic and entropy computations required significantly more memory than minimization calculations. Additional computational efficiency was achieved by separating the memory intensive computations from low-memory calculations. Each complex typically took about 2-3 days on NYU Abu Dhabi’s Linux clusters. We have previously reported some of above computational details and protocols^23; 24^.

## Abbreviations used

COVID-19: coronavirus disease 2019
SARS-CoV-2: severe acute respiratory syndrome coronavirus 2
ACE2: angiotensin-converting enzyme 2
RBD: receptor-binding domain
RBM: receptor-binding motif
S: SARS-CoV-2 Spike protein

## Acknowledgments

This research was carried out on the High Performance Computing resources at New York University Abu Dhabi. We thank Neville Sanjana for his valuable comments and suggestions.

## Author contributions

HHG and KCG conceived the project and wrote the paper. HHG carried out structural modeling computations; BM worked on code optimization; and AT developed the variant calling algorithm and performed analysis of Spike variants. All authors contributed to providing critical comments and editing the paper.

---

**Supplementary Figure 1:**
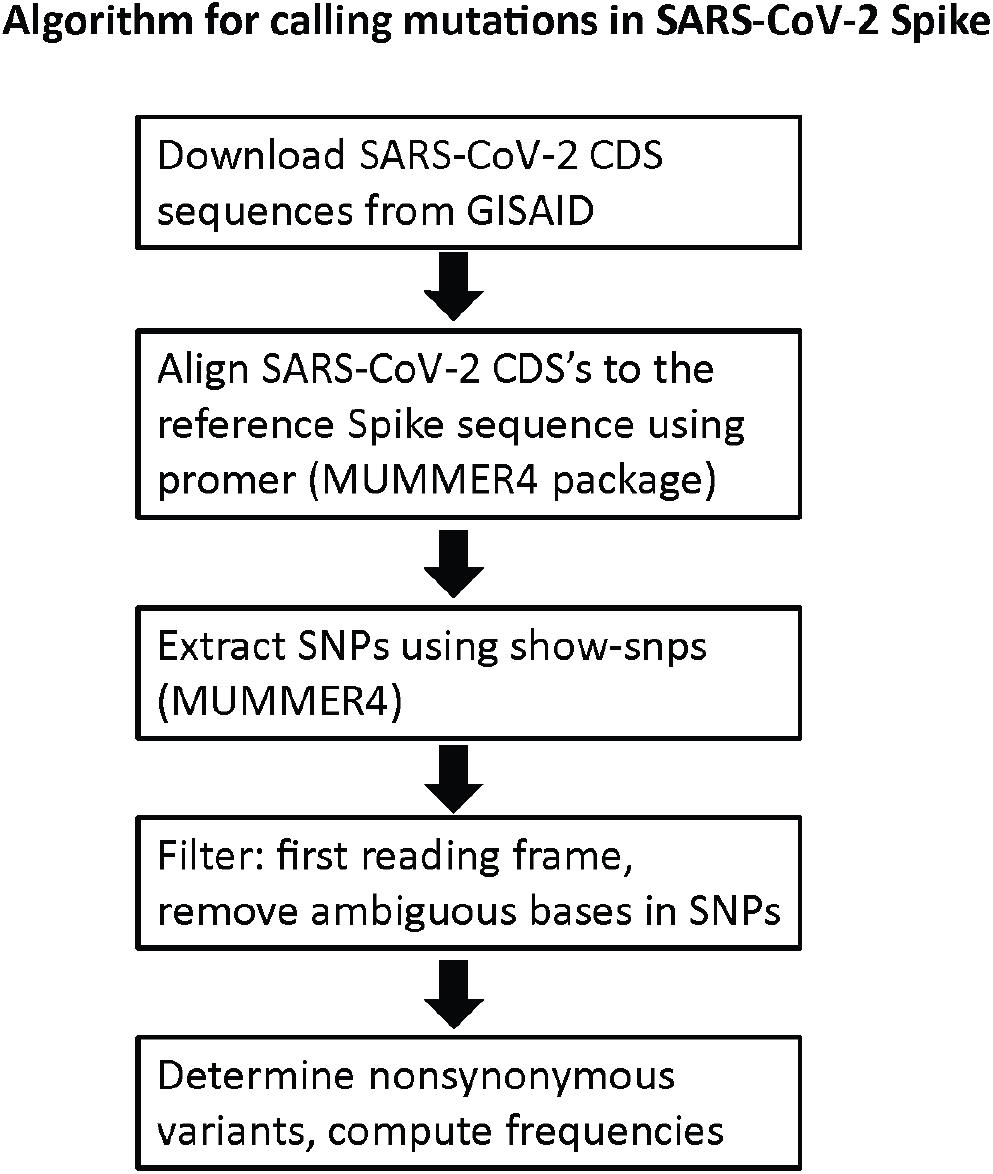
Computational pipeline for calling mutations in SARS-CoV-2 Spike. Available at GitHub (https://github.com/GunsalusPiano/covid-snp-analysis).

**Supplementary Figure 2:**
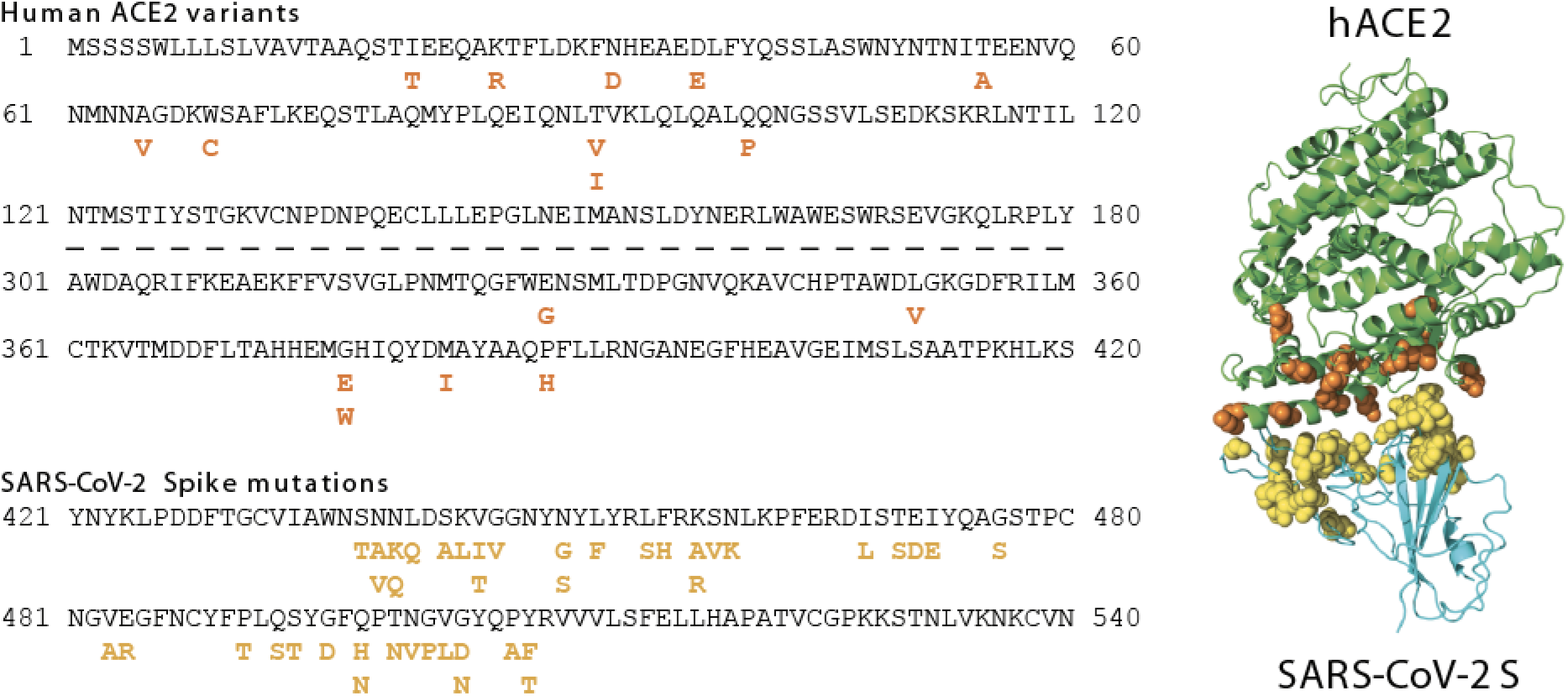
Sequence and structural contexts of Spike mutations (magenta) and hACE2 variants (orange) at the binding interface.

**Supplementary Figure 3:**
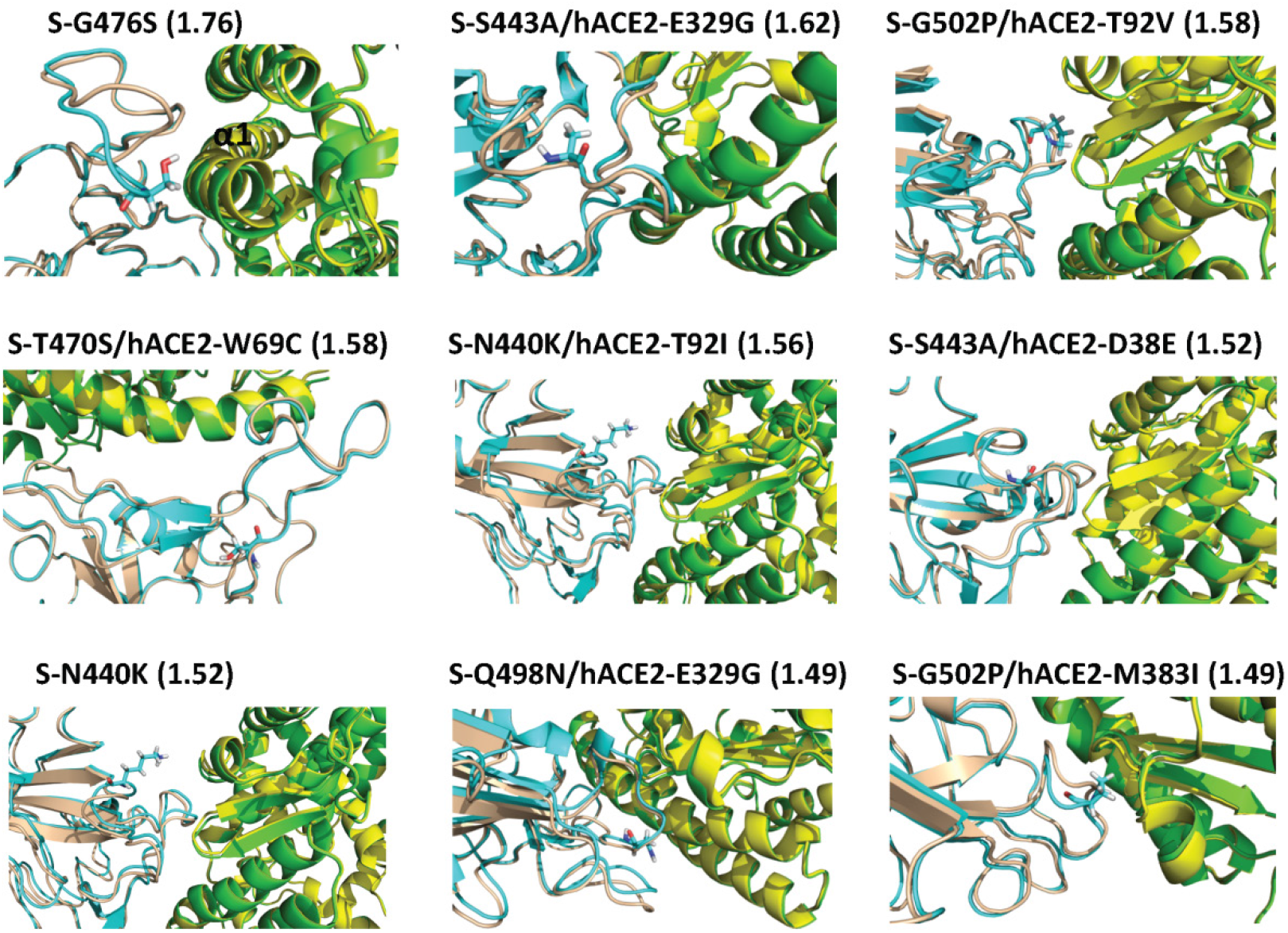
Conformational changes at the S-hACE2 interface induced by the top 9 high-affinity mutations, ordered by normalized affinity values. Color scheme: WT complex, hACE2 in yellow and S in wheat; mutant complexes, hACE2 in green and S in cyan.

